# Momo – Multi-Objective Metabolic mixed integer Optimization: application to yeast strain engineering

**DOI:** 10.1101/476689

**Authors:** Ricardo Andrade, Mahdi Doostmohammadi, João L. Santos, Marie-France Sagot, Nuno P. Mira, Susana Vinga

## Abstract

In this paper, we explore the concept of multi-objective optimization in the field of metabolic engineering when both continuous and integer decision variables are involved in the model. In particular, we propose a multi-objective model that may be used to suggest reaction deletions that maximize and/or minimize several functions simultaneously. The applications may include, among others, the concurrent maximization of a bioproduct and of biomass, or maximization of a bioproduct while minimizing the formation of a given by-product, two common requirements in microbial metabolic engineering.

Production of ethanol by the widely used cell factory *Saccharomyces cerevisiae* was adopted as a case study to demonstrate the usefulness of the proposed approach in identifying genetic manipulations that improve productivity and yield of this economically highly relevant bioproduct. We did an *in vivo* validation and we could show that some of the predicted deletions exhibit increased ethanol levels in comparison with the wild-type strain. The multi-objective programming framework we developed, called Momo, is open-source and uses PolySCIP^‡^ as underlying multi-objective solver. Momo is available at http://momo-sysbio.gforge.inria.fr

## 1 Introduction

Multi-objective (multi-criteria) optimization is a method used to tackle problems when several objective functions have to be optimized simultaneously. Particularly, multi-objective programming is a class of optimization problems that have more than one objective function over a set of constraints. Except under limited circumstances, it is not possible to find a single optimal solution which simultaneously optimizes all of the objective functions, *i.e.*, in general, the resolution of such problems leads to a trade-off curve (known as “Pareto frontier”) such that at any point of the curve, a compromise is made in favor of some objective function.

Multi-objective optimization has been extensively exploited in science and engineering. It has become a powerful tool for solving real-world problems and is also used in numerous bioinformatics and computational biology as shown in [1]. Successful engineering of microbial catalysts often needs the optimization of more than one feature, for example, the maximization of the production of a desired product while minimizing the synthesis of a by-product or maximizing the production of a product that is an end-point metabolite and therefore competes with growth for the carbon source. The most common algorithms used to guide strain design do not focus on a multi-objective approach but instead, in general, work within a bi-level framework (OptKnock is the more paradigmatic case [2, 3]), adopt a max-min approach (*e.g.* RobustKnock [4]), or exploit a consensus framework (*e.g.* the recently described OptPipe method [5]), as recently reviewed by [6].

There are also some other more generic approaches such as the one of Hädicke and Klamt [7] that is based on constrained minimal cut sets that allow disabling or preserving some cell functionalities when computing a gene deletion strategy. This method is generic and can be used to emulate other approaches such as OptKnock, but it does not handle the optimization of multiple simultaneous compounds.

The use of multi-objective optimization for strain engineering is usually performed exploring heuristic/meta-heuristic approaches. Examples of these studies are the work of Maia *et al.* [8], in which the authors model the problem of identifying deletions to increase production of a compound of interest using multi-objective optimization, and then use an evolutionary algorithm to obtain the solutions. The developed approach was validated by improving the capability of *Escherichia coli* to produce succinic acid. Another example of studies exploring multi-objective optimization by heuristic methods is the work of Costanza *et al.* [9] that uses such an approach to seek the deletions that may optimize multiple cellular functions simultaneously, thereby allowing what the authors called a genetic design of the strains.

Although heuristic methods are usually computationally less expensive than exact ones, they do not guarantee that an optimal solution is found. In this work, we present an exact integer-linear multi-objective optimization methodology that can be explored for strain engineering, thereby expanding the current set of tools available for this objective. In our framework, called Momo given a metabolic network and a set of target reactions, the user can identify possible reaction deletions that could improve the flux of those targets, always in a multi-objective setting. To the best of our knowledge, this is the first time that such a tool has been developed combining these features.

To validate the approach, Momo was used to model ethanol production undertaken by *S. cerevisiae* cells, and deletion of reactions that could optimize ethanol production were predicted and tested *in vivo*. Simultaneous maximization of growth and ethanol were used as objective functions. Despite many years of utilization in the industrial biotechnology sector, alcoholic fermentation undertaken by yeast cells gained a further boost in recent years with an increase in the interest of exploring ethanol as a biofuel [10].

Previous works have also aimed at exploring non-linear multi-objective optimization formulations to model *Saccharomyces cerevisiae* production of ethanol [11, 12]. Although these two approaches provided some understanding concerning the control and optimization of the small metabolic system used, no strategies to improve ethanol production could be designed from the results obtained.

We start by presenting in Section 2 an overview of metabolic network modeling and of the most used method for metabolic engineering, namely OptKnock that adopts a bi-level model (Section 2.1 and Section 2.2). In Section 2.3, we introduce a bi-objective approach that will form the core of the Momo method we developed, together with a classical enumeration technique that is required when there is more than one optimal solution and all need to be listed. In Section 2.4, we describe the experimental context in which the yeast fermentations were made. In Section 3, we present the experimental results obtained *in silico*, and the experimental validation that was performed. Finally, in Section 4 we discuss such results and present some of the perspectives of this work. The Appendix S1 provides additional theoretical material on multi-objective optimization.

## 2 Methods and models

In this section, we describe some of basic aspects related to modeling metabolic networks in steady-state conditions and briefly describe the well-known algorithm OptKnock, this being an important feature to better understand the modelling behind Momo which was based on this. We also present the model of Momo and its implementation aspects as well as some details about the fermentations performed to validate our method.

### 2.1 Modeling and optimization of metabolic networks

Genome-scale metabolic models are linked with conservation of mass equations for each metabolite in the metabolic network. The quantitative metabolic capabilities of a given organism can therefore be calculated as follows. Let *I* = 1*,…, m* index the metabolites and *J* = 1*,…, n* index the reactions of a given metabolic network. Assume that *x_i_* denotes the concentration of metabolite *i*, and *v_j_* denotes the flux of reaction *j*. The system dynamics is then described by the following equations:

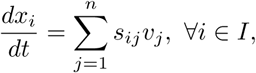

where *s_ij_* are the stoichiometric coefficients of metabolite *i* in reaction *j*. Observe that the stoichiometric coefficients *s_ij_* can be combined into the so-called stoichiometric matrix *S* = [*s_ij_*]_*m×n*_, where each row corresponds to a metabolite and each column to a reaction. In this paper, we consider the steady-state approximation in which the metabolite concentrations are assumed to be constant. The above equation therefore reduces to:

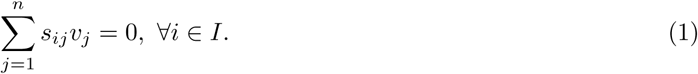

Since the number of reactions (denoted by *n*) is higher that the number of metabolites (denoted by *m*) in the majority of the real-world metabolic networks, there can be an infinite number of reaction fluxes satisfying the system of equations (1). One therefore needs to select from the set of feasible solutions only those that ensure cell viability and, in this context, one option currently applied is to maximize the flux of the biomass reaction (*v_biomass_*). This is obtained by solving a continuous linear programming problem. In order to formulate such a problem, assume that *LB_j_* and *U B_j_* denote the given lower bound and upper bound on the flux of reaction *j*, respectively. This optimization problem can therefore be written as:

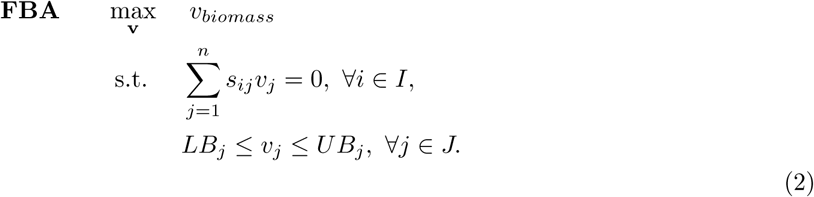

The above mathematical program is an essential part of Flux Balance Analysis (FBA) [13–"19] which plays a key role in optimizing cell metabolism and enables us to investigate both the theoretical and operative capabilities of such metabolism.

### 2.2 Bi-level model

We now describe the mathematical formulation of the OptKnock [2] procedure in detail. OptKnock was developed to identify multiple reaction deletions that may lead to the overproduction of a target chemical while cellular growth is maximized. This hierarchical optimization model comprises two decision or competing strategies, namely a cellular objective and a chemical production objective. It therefore is formulated as a Bi-Level Mixed Integer Program (BLMIP). Two binary variables, denoted by *y_j_, j ϵ J*, are defined which enable to perform reaction removals inside the model by multiplying both *LB_j_* and *U B_j_* by (1 *− y_j_*) in the lower bound and upper bound constraints as follows:

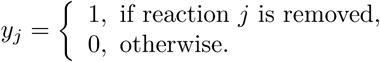

Assume that *K* denotes the given number of knock-outs. Then the Bi-Level Mixed Binary (linear) Program (BLMBP), which is a hierarchy of two optimization problems, can be mathematically formulated as follows:

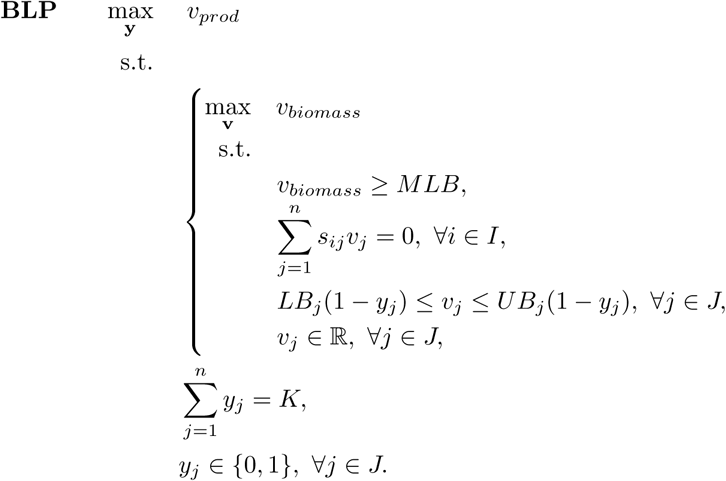

where *v_prod_* corresponds to the flux of a chemical of interest, the vectors **v** and **y** denote (*v*_1_*,…, v_n_*) and (*y*_1_*,…, y_n_*) respectively, and *M BL* represents a minimum level of biomass production. We may also need to fix a substrate uptake that in this model we suppose is specified through the vectors *LB* and *U B*.

There is also one variation of OptKnock, known as SimpleOptKnock and implemented in the COBRA toolbox [20] that does not rely on a bi-level model, but on solving many linear programming problems.

This method consists of solving *J* instances of a variant of (2). For each reaction, its upper and lower flux bounds are fixed to zero. Under these new constraints that simulate the deletion of that reaction, an optimization of the biomass is performed. The obtained biomass value (maximum) is then fixed (upper and lower flux bounds) and this new model is subsequently optimized for the bioproduct of interest. The algorithm output, as implemented in the COBRA Toolbox, reports, for each reaction knock-out, the maximum biomass and the maximum and minimum values for the target compound.

### 2.3 Momo method

We now describe the Momo method developed in this study. This is based on a bi-objective model, whose characteristics are presented in the next subsection, and on an enumeration technique that is discussed just afterwards. Finally, Momo itself, some of its implementation details and a short comparison with OptKnock is discussed in the last part of this section.

#### 2.3.1 Bi-objective model

In this section, we propose and describe a novel formulation for predicting reaction removal strategies in order to overproduce a target chemical in metabolic networks.

The formulation is based on multi-objective programming, and could be used for more than two objectives. In the case-study of this paper, we however concentrate on only two. Those are growth on one hand, and target chemical overproduction on the other, knowing that in general the two compete inside a cell, *i.e.*, producing more of a target chemical leads to less production of biomass. We use such formulation to propose reaction knock-outs in order to reach the second objective while achieving an acceptable level of the first, that is, of cell growth rate. A detailed description of multi-objective optimization (definitions and algorithms) is presented in the Appendix, along with the main references.

We briefly describe however here one key concept, namely of the Pareto-efficient frontier, whose precise definition is given in the Appendix. Informally speaking, the Pareto-efficient frontier represents a set of solution points that are optimal in the sense that we cannot improve one objective without worsening another one. One example of a Pareto-optimal frontier is given in Fig. 1, where the points A, B, C, D, and E are considered Pareto-optimal.

**Fig 1.**
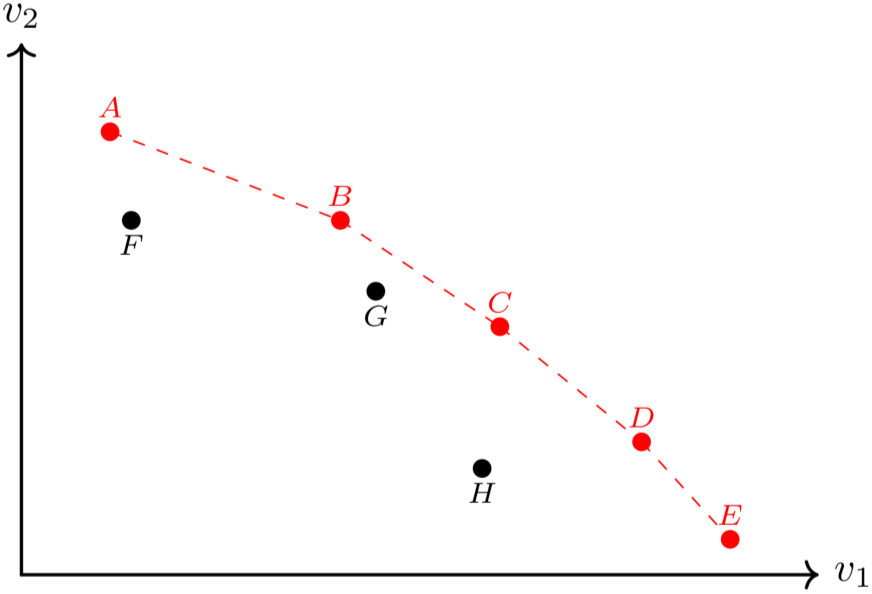
An example of a Pareto-efficient frontier with two objectives, *v*_1_ and *v*_2_. The red dots A, B, C, D, and E are examples of optimal choices while the points F, G, and H represent non-optimal solutions since we can improve one of the objectives without worsening the other.

Considering the notations of Section 2.1, we partition the set *J* into (*J_E_, J_I_*) in such a way that *J_E_* indexes the *essential reactions* which are required for the cell to meet the biological objectives and *J_I_* indexes the *inessential reactions* that are not necessary for the cell. The Bi-Objective Mixed Binary (linear) Program (BOMBP) can therefore be mathematically formulated as follows:

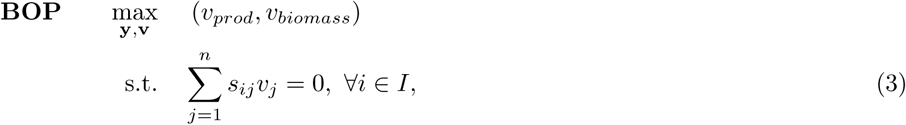

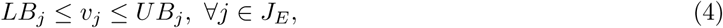

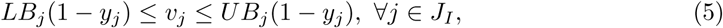

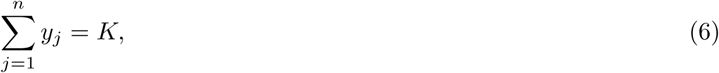

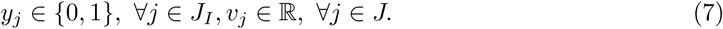

Notice that in BOP, the binary variables *y_j_* are only attributed to inessential reactions. We highlight that instead of formulating the problem of identifying reaction knock-out strategies with a bi-level model which gives exactly one optimal solution, we formulate this problem using a bi-objective model (as presented above) which provides a number of optimal solutions on the trade-off Pareto-optimal frontier. Hence, we are able to analyze each optimal solution of the bi-objective model more carefully in order to find the most suitable reaction deletion scenarios.

We describe next the technique used to enumerate all optimal solutions.

#### 2.3.2 Enumeration

The models presented so far have the limitation of predicting only one set of *K* deletions per point in the Pareto frontier. Since we may have many sets of deletions leading to the same production rates, it would be interesting to enumerate all sets of possible deletions for each point in the Pareto frontier.

We therefore now describe a classical enumeration technique that can be applied in conjunction with the models presented so far in order to enumerate all equivalent deletions, that is, all deletion sets that allow the same production rates of the desired product and of biomass. We also present a modification of this technique in order to enumerate not all equivalent deletions, but a subset of these deletions that allows a flux distribution with a minimal change of the fluxes in comparison to a reference flux vector. In our case, we use as reference the flux distribution obtained by the FBA model since it represents the one of the wild-type organism. This idea of minimizing the distance between a reference flux distribution and the mutant flux distribution is known as minimization of metabolic adjustment (Moma) [21].

Let *Z\** be set of points of the Pareto frontier obtained by solving the BOP. For each 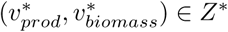, we construct the following feasibility problem:

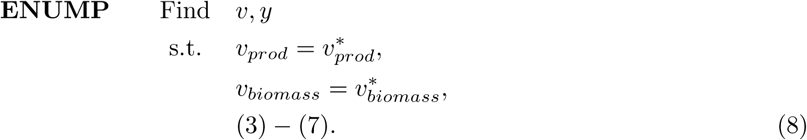

Let 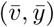 be one solution of ENUMP. The vector 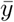 represents one set of deletions. Let *ρ*(*y*) be a function that takes a vector of integers and returns the set of indexes *L* such that for all *l ϵ L, y_l_* = 1. Consider the following constraint:

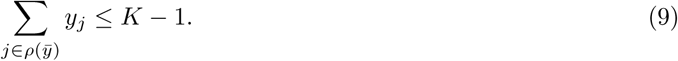

Adding Equation (9) to ENUMP will exclude the set defined by 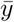 from the set of solutions and also any other set that contains it. Solving the problem again will either give a new deletion set, or a certificate proving that no other equivalent solution exists. Iterating over these steps allows to enumerate all sets of equivalent deletions for each point in the Pareto frontier.

Observe that the amount of sets with equivalent deletions may grow considerably if *K* is large. One alternative is to enumerate only a subset of the deletions. For that we can add an objective function to ENUMP so that only those deletion sets that are closer to a reference flux set are enumerated. One possible objective function would be to consider the strategy introduced in [21], minimizing the distance between the wild-type and the mutant distribution through quadratic programming. Here we considered a similar idea using the Manhattan norm instead of the Euclidean one. This decision allows us to stay in a integer-linear model and also allows sparser solutions.

We can now define the new problem for the enumeration procedure as follows:

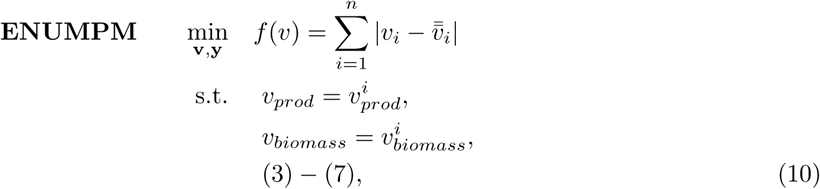

where 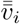 is the optimal solution obtained from FBA. The norm on the objective function of ENUMPM can be modeled using a classical reformulation technique where *f* (*v*) can be reformulated as:

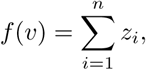

where:

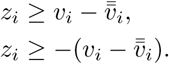

For the enumeration, it suffices to add two constraints. The first is a constraint fixing the value of the objective function to the optimal one obtained by solving ENUMPM for the first time:

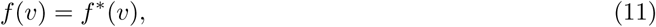

where *f*\* (*v*) is the optimal value of ENUMPM. Then at each step, one should also add Equation (9) as explained previously.

Using ENUMPM in the enumeration scheme presented here allows to obtain all deletion sets that enable the desired production rates, and, at the same time, have minimum difference in the flux distribution compared to the wild-type theoretical fluxes.

### 2.3.3 Implementation of Momo

The ideas presented in the previous sections (Section 2.3.1 Section 2.3.2) were implemented in the software Momo. The latter is coded in C++ and uses SCIP [22] and PolySCIP [23] as solvers for the optimization models.

Momo allows to solve instances of the FBA, BOP, ENUMP and ENUMPM models, using exact methods, and further allows for variations of those when there are more than two objective functions, which may include for instance maximizing and/or minimizing other bio-products. The software is open-source and available at http://momo-sysbio.gforge.inria.fr/.

We present below a sequence of steps that can be used with Momo to predict knockout strategies for the overproduction of a desired product:

1. Solve an instance of FBA in order to obtain a reference flux distribution.
2. Identify the reactions that have a non-zero flux in the previous step and consider them as knock-out candidates.
3. Solve an instance of the BOP or of one of the multi-objective models to obtain points in the Pareto curve.
4. For each point, fix the values of the objective functions and enumerate all the knock-outs using ENUMP or ENUMPM.

#### 2.4 Yeast fermentations

To biologically evaluate the proposed approach, we applied Momo with the objective of finding gene deletions that could optimize the production of ethanol while also maximizing biomass in yeast.

The fermentations were performed using the wild-type strain *Saccharomyces cerevisiae* BY4741 (MATa, his3∆1, leu2∆0, met15∆0, ura3∆0) and W303 (MAT*α* ade2-1, his3-1,15 leu2-3,112 trp1-1 ura3-1). BY4741 was acquired from the Euroscarf collection while the W303 strain was kindly provided by Dr Dennis Winge (University of Utah). The mutant strains devoid of genes *FUM1*, *NDI1*, *MDH1*, *ADE3*, *ALT1*, *PGI1*, *FUM1*, *SCS7 ADO1*, *BAT2*, *LOT6*, *IDP1*, *LOT6*, *YND1*, *FRD1*, *IPT1* and *SUR2* were derived from the BY4741 background (and were also acquired from the Euroscarf collection), while strains devoid of *SDH2* or of *SDH6* and *SDH7* were kindly provided by Dr Dennis Winge (University of Utah).

For the very high gravity fermentations, a protocol similar to the one described in [24] was used. Briefly, the strains were cultivated over-night at 30°C in YPD growth medium (20 g/L glucose, 20 g/L bactopeptone and 10 g/L yeast extract), after which they were inoculated in the fermentation medium YPF which differs from YPD by having 300 g/L glucose as well as 240 mg/L leucine, 80 mg/L histidine, 80 mg/L methionine and 80 mg/L uracil which were added to the medium to complement the auxotrophies of the BY4741 strain. The different fermentations were assessed measuring cell growth (through the increase in culture OD_600nm_) and accompanying the alterations in the concentration of glucose, ethanol and glycerol in the supernatant. For this, 10*µ*L of each supernatant sample were separated in an Aminex HPX-87 H Ion Exchange Chromatography column, eluted at room temperature with 0.005 M H2SO4 at a flow-rate of 0.6 mL min-1 during 30 minutes, using a refractive-index detector. Under such experimental conditions, glucose had a retention time of 8.3 minutes and ethanol of 19.4 minutes. Reproducibility and linearity of the method were tested, and the concentrations were estimated based on appropriate calibration curves.

## 3 Results

### 3.1 Multi-objective optimization-based modelling of ethanol production in *S. cerevisiae*

To simulate the yeast-based production of ethanol, the Yeast 5.01 model [25] was used constraining the flux of glucose uptake (`r_1714`, D-glucose exchange, lower-bound limited to −10), the electron transfer to O2 (corresponding to reaction `r_0438`, ‘ferrocytochrome-c:oxygen oxidoreductase’, upper-bound limited to 10) and the isomerization reaction between dihydroxyacetone phosphate and glyceraldeyde-3-phosphate (corresponding to reaction `r_1054`, ‘triose-phosphate isomerase’, upper-bound limited to 6). The constraint of the oxygen reduction reaction was necessary to predict ethanol production as otherwise all the glucose was channeled for biomass growth. Constraining the isomerization reaction was performed in order to predict the co-production of ethanol and glycerol, something that the non-modified Yeast 5.01 model could not do but that is well described to occur during yeast alcoholic fermentation due to the need of recycling NADH. Also, all the infinite bounds in the Yeast 5.01 model, -INF and +INF, were set to −1000 and +1000, respectively.

Under the established conditions and using biomass as the objective function, the predicted flux of ethanol was 8.8 mmol/g dry weight/h, while the predicted flux for glycerol production was of 2.2 mmol/g dry weight/h. When changing the objective function towards “maximization of ethanol production”, no flux for the biomass could be predicted, thus reflecting the competing nature that exists between ethanol production and growth (Fig. 2).

**Fig 2.**
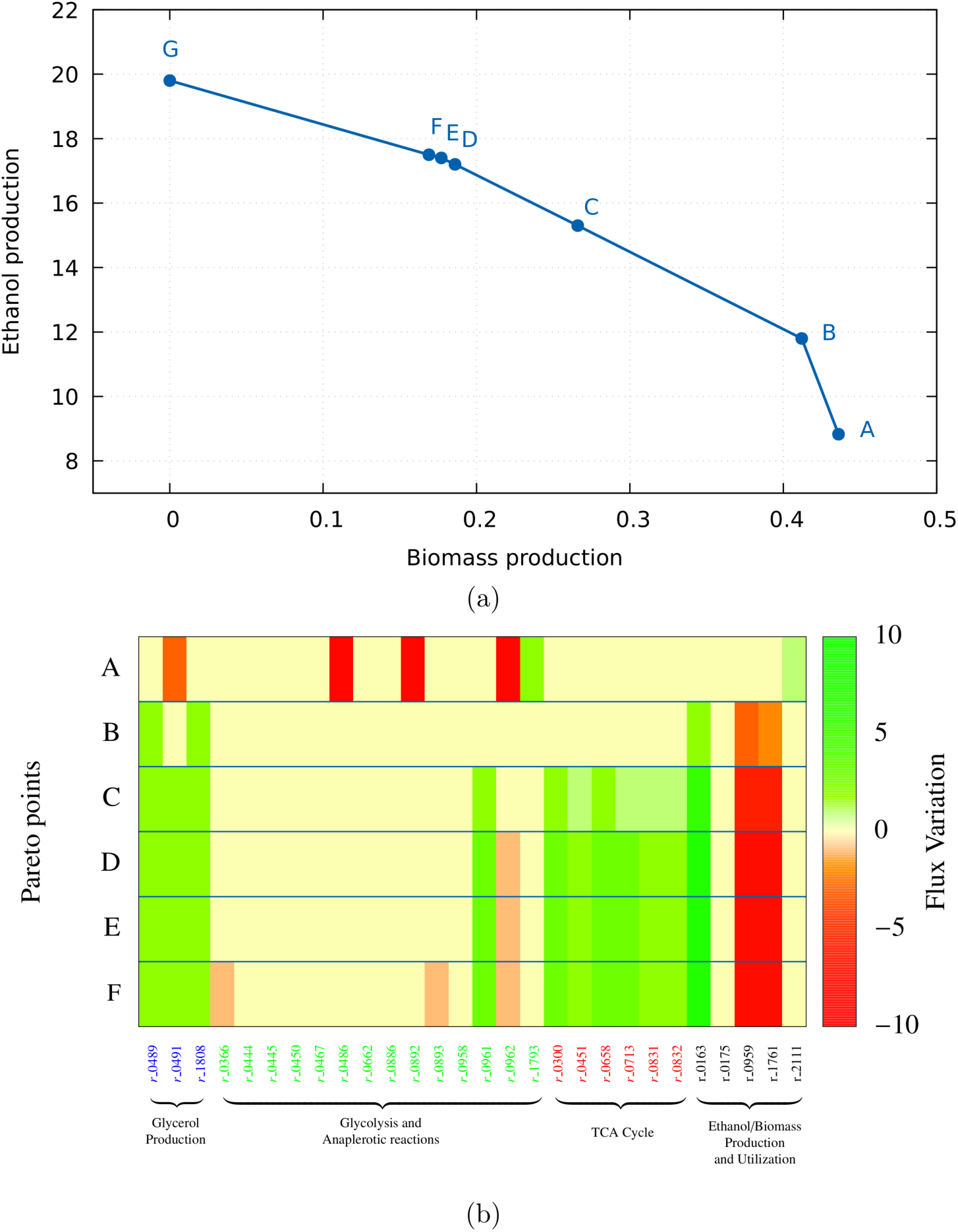
(a) Pareto-optimal points obtained upon simulation of yeast alcoholic fermentation maximizing biomass and ethanol production. The fluxes obtained for ethanol and biomass production on the A-G points of the Pareto frontier are as follows: A(8.8;0.43); B(11.8;0.41);C(15.3;0.27);D(17.2;0.19);E(17.4;0.18); F(17.5;0.17); G(19.8;0.0). (b) Heatmap illustrating the difference of the fluxes between the wild-type and mutants corresponding to each Pareto point. Considering reactions related to glycerol production, glycolysis, TCA cycle, and ethanol production and utilization.

Simultaneous maximization of ethanol production and biomass growth using Momo resulted in the identification of 6 Pareto frontier points (Fig. 2-(a)).

Point A predicts fluxes of ethanol and biomass growth identical to those predicted when biomass is used as the objective function (Fig. 2-(a)) (8.83 and 0.44 respectively). Along the Pareto frontier, an increased amount of ethanol is predicted, achieving a flux of 17.5 (Fig. 2-(a)) while the flux of the predicted biomass is about 0.18 (in fact the maximum predicted flux for ethanol is of 18.8 but with no biomass prediction).

It is clear from the analysis of the data shown in Fig. 2 that the maximization of ethanol observed along the Pareto frontier is accompanied by a significant decrease in biomass flux, with the exception of the point B in which ethanol flux increases to 11.8 but biomass growth decreases minimally compared with the results that are obtained when biomass is the sole objective function (Figs 2-(a) and 2-(b)).

A closer examination of the distribution of the fluxes obtained upon implementation of the multi-objective setting for the different points of the Pareto frontier (in part shown in Fig. 2-(b) and further detailed in Table S1) shows that the predicted increase in ethanol production is, at first, obtained at the expense of eliminating glycerol production (whose associated reaction fluxes drop to zero) and of reducing the activity of various reactions that consume pyruvate, including pyruvate dehydrogenase and pyruvate carboxylase (Fig. 2-(b)). On the other hand, the glycolytic reactions involved in pyruvate replenishing are predicted to increase their fluxes along the different Pareto frontier points as well as pyruvate decarboxylase and alcohol dehydrogenase, the enzymes driving ethanol synthesis from pyruvate (Fig 2-(b)). The reactions diverting acetaldehyde from ethanol synthesis show a reduced flux, as do the reactions involved in ethanol consumption (Fig 2-(b)).

Another important observation concerns the reduction in the flux of various enzymes involved in the TCA cycle and in amino acid biosynthesis, something that could be attributed to the reduction that is observed in biomass, this effect being much more marked for the last points of the Pareto frontier where the decrease in biomass is more evident (Fig. 2-(b)). Overall, it can be seen that the distribution of fluxes around the yeast metabolic network during alcoholic fermentation in the bi-objective setting is first performed at the expense of reducing the formation of byproducts but not growth.

### 3.2 Identification of deletions leading to improved ethanol production

The use of an *in silico* simulation to identify gene deletions is one of the most widely adopted strategies to improve the microbe-based production of desired compounds. In this direction, we used Momo to identify putative deletions that could lead to an improved ethanol production undertaken by yeast cells. For this, a set of possible deletions was generated for each of the points of the Pareto frontier (see Table S2). This was accomplished by fixing the objective functions *v_ethanol_* and *v_biomass_* to the value of each point and then using the Momo enumeration technique (as explained in Section 2.3.2 and in Step 4 of Section 2.3.3).

A selected set of the results obtained is shown in Table 1 while the full list of deletions is shown in Table S2. The highest number of candidate reactions for deletion was obtained for point G (317), which is expected since this is the point where no biomass is predicted therefore giving a higher flexibility for identification of candidate reaction deletions. A closer look into the sets of solutions obtained for all Pareto frontier points revealed possibilities for deleting reactions associated with the metabolism of glycerol, pyruvate (*e.g.* pyruvate dehydrogenase or acetaldeyde dehydrogenase), with the activity of Krebs cycle (e.g. malate dehydrogenase, fumarase, pyruvate dehydrogenase, oxoglutarate dehydrogenase), with the respiratory chain (*e.g.* succinate dehydrogenase, ferrocytochrome-c:oxygen oxidoreductase), with the biosynthesis of amino acids, with the metabolism of complex lipids including ergosterol, sphingolipids or ceramides or with the synthesis of glutathione.

**Table 1.**
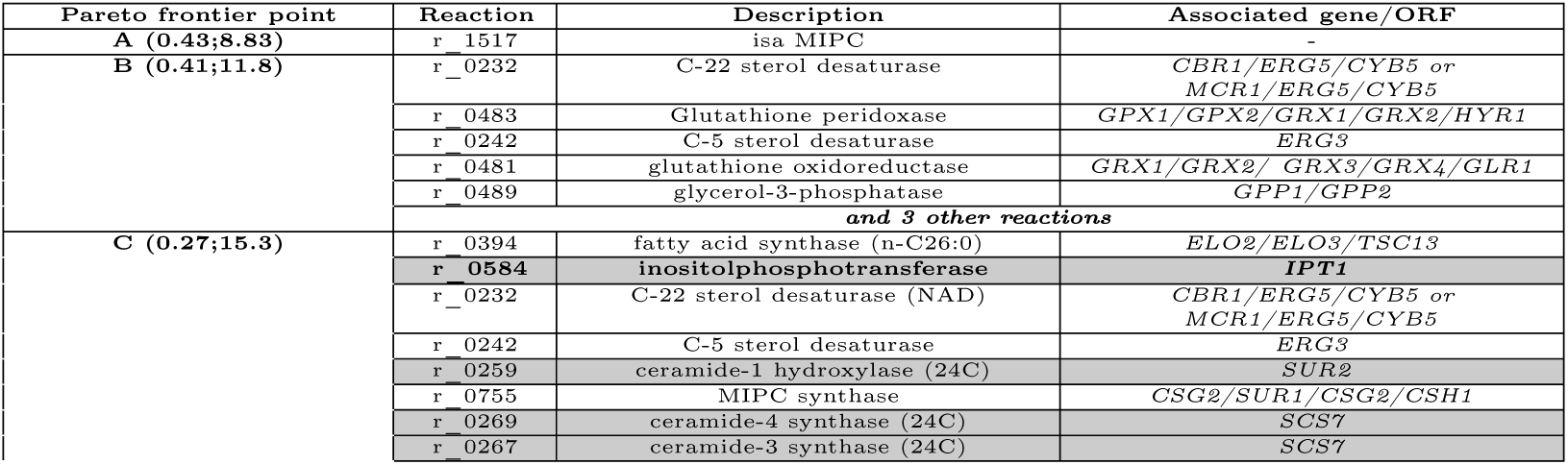
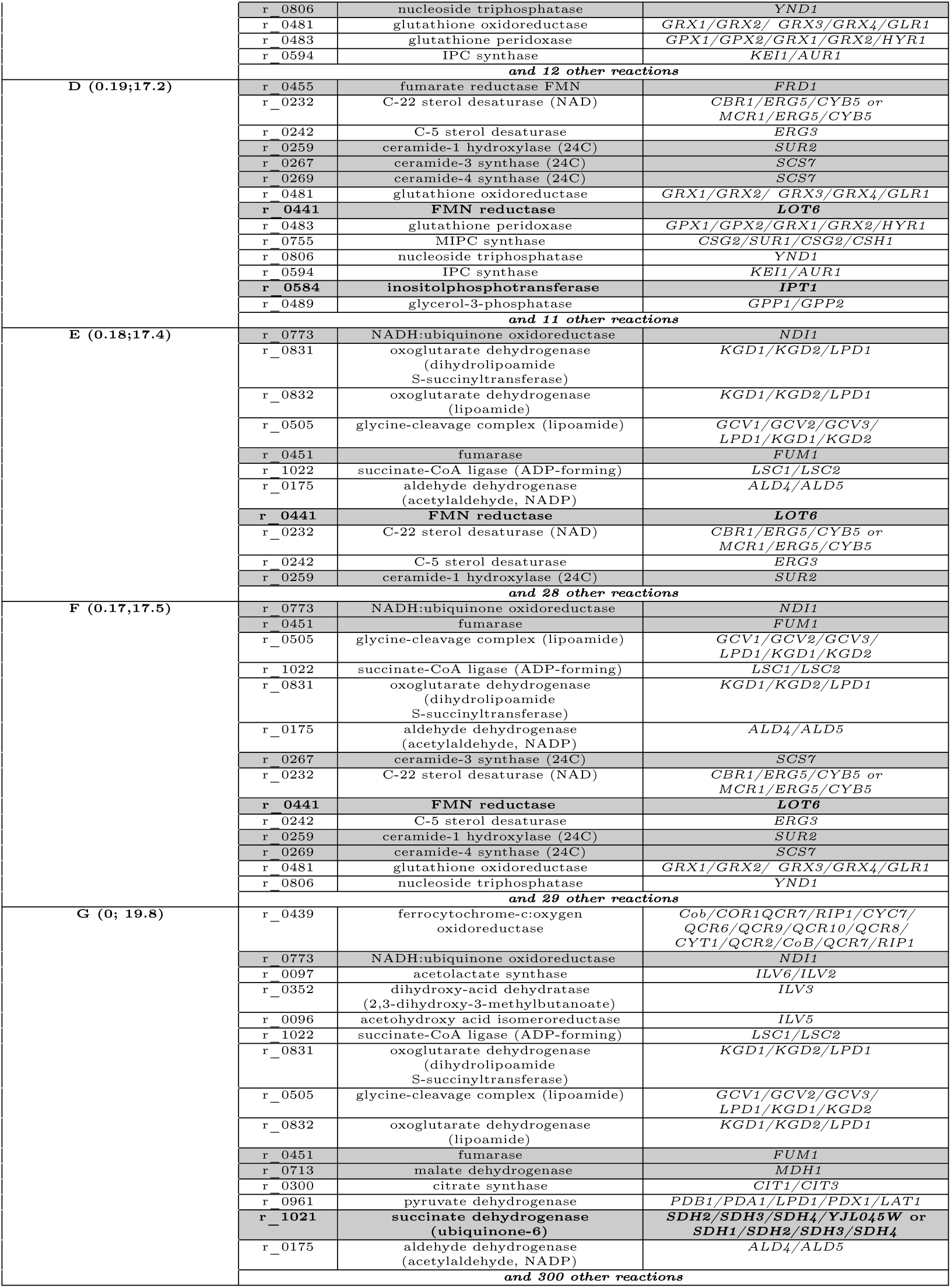
Sub-set of the deletions identified by Momo for each point of the Pareto frontier shown in Fig 2. The complete list of reactions identified by Momo is provided in Table S2. The reaction number, the corresponding function and the associated *S. cerevisae* genes are indicated. The flux of biomass and ethanol predicted for each point of the Pareto frontier is indicated in brackets. A selected set of 10 deletions tested *in vivo* are highlighted in gray and those deletions leading to improved production are further highlighted in bold.

| Pareto frontier point | Reaction | Description | Associated gene/ORF |
| --- | --- | --- | --- |
| **A (0.43;8.83)** | r_1517 | isa MIPC |  |
| **B (0.41;11.8)** | r_0232 | C-22 sterol desaturase | *CBR1/ERG5/CYB5* or *MCR1/ERG5/CYB5* |
|  | r_0483 | Glutathione peroxidase | *GPX1/GPX2/GRX1/GRX2/HYR1* |
|  | r_0242 | C-5 sterol desaturase | *ERG3* |
|  | r_0481 | glutathione oxidoreductase | *GRX1/GRX2/GRX3/GRX4/GLR1* |
|  | r_0489 | glycerol-3-phosphatase | *GPP1/GPP2* |
|  | **and 3 other reactions** |  |  |
| **C (0.27;15.3)** | r_0394 | fatty acid synthase (n-C26:0) | *ELO2/ELO3/TSC13* |
|  | **r_0584** | **inositolphosphotransferase** | ***IPT1*** |
|  | r_0232 | C-22 sterol desaturase (NAD) | *CBR1/ERG5/CYB5* or *MCR1/ERG5/CYB5* |
|  | r_0242 | C-5 sterol desaturase | *ERG3* |
|  | r_0259 | ceramide-1 hydroxylase (24C) | *SUR2* |
|  | r_0755 | MIPC synthase | *CSG2/SUR1/CSG2/CSH1* |
|  | r_0269 | ceramide-4 synthase (24C) | *SCS7* |
|  | r_0267 | ceramide-3 synthase (24C) | *SCS7* |

**Table 1.** Sub-set of the deletions identified by MOMO for each point of the Pareto frontier shown in Fig 2.
|  |  |  |  |
| --- | --- | --- | --- |
|  | r_0806 | nucleoside triphosphatase | YND1 |
|  | r_0481 | glutathione oxidoreductase | GRX1/GRX2/ GRX3/GRX4/GLR1 |
|  | r_0483 | glutathione peroxidase | GPX1/GPX2/GRX1/GRX2/HYR1 |
|  | r_0594 | IPC synthase | KEI1/AUR1 |
|  | **and 12 other reactions** |  |  |
| D (0.19;17.2) | r_0455 | fumarate reductase FMN | FRD1 |
|  | r_0232 | C-22 sterol desaturase (NAD) | CBR1/ERG5/CYB5 or MCR1/ERG5/CYB5 |
|  | r_0242 | C-5 sterol desaturase | ERG3 |
|  | r_0259 | ceramide-1 hydroxylase (24C) | SUR2 |
|  | r_0267 | ceramide-3 synthase (24C) | SCS7 |
|  | r_0269 | ceramide-4 synthase (24C) | SCS7 |
|  | r_0481 | glutathione oxidoreductase | GRX1/GRX2/ GRX3/GRX4/GLR1 |
|  | r_0441 | **FMN reductase** | **LOT6** |
|  | r_0483 | glutathione peroxidase | GPX1/GPX2/GRX1/GRX2/HYR1 |
|  | r_0755 | MIPC synthase | CSG2/SUR1/CSG2/CSH1 |
|  | r_0806 | nucleoside triphosphatase | YND1 |
|  | r_0594 | IPC synthase | KEI1/AUR1 |
|  | r_0584 | **inositolphosphotransferase** | **IPT1** |
|  | r_0489 | glycerol-3-phosphatase | GPP1/GPP2 |
|  | **and 11 other reactions** |  |  |
| E (0.18;17.4) | r_0773 | NADH:ubiquinone oxidoreductase | NDI1 |
|  | r_0831 | oxoglutarate dehydrogenase (dihydrolipoamide S-succinyltransferase) | KGD1/KGD2/LPD1 |
|  | r_0832 | oxoglutarate dehydrogenase (lipoamide) | KGD1/KGD2/LPD1 |
|  | r_0505 | glycine-cleavage complex (lipoamide) | GCV1/GCV2/GCV3/ LPD1/KGD1/KGD2 |
|  | r_0451 | fumarase | FUM1 |
|  | r_1022 | succinate-CoA ligase (ADP-forming) | LSC1/LSC2 |
|  | r_0175 | aldehyde dehydrogenase (acetylaldehyde, NADP) | ALD4/ALD5 |
|  | r_0441 | **FMN reductase** | **LOT6** |
|  | r_0232 | C-22 sterol desaturase (NAD) | CBR1/ERG5/CYB5 or MCR1/ERG5/CYB5 |
|  | r_0242 | C-5 sterol desaturase | ERG3 |
|  | r_0259 | ceramide-1 hydroxylase (24C) | SUR2 |
|  | **and 28 other reactions** |  |  |
| F (0.17;17.5) | r_0773 | NADH:ubiquinone oxidoreductase | NDI1 |
|  | r_0451 | fumarase | FUM1 |
|  | r_0505 | glycine-cleavage complex (lipoamide) | GCV1/GCV2/GCV3/ LPD1/KGD1/KGD2 |
|  | r_1022 | succinate-CoA ligase (ADP-forming) | LSC1/LSC2 |
|  | r_0831 | oxoglutarate dehydrogenase (dihydrolipoamide S-succinyltransferase) | KGD1/KGD2/LPD1 |
|  | r_0175 | aldehyde dehydrogenase (acetylaldehyde, NADP) | ALD4/ALD5 |
|  | r_0267 | ceramide-3 synthase (24C) | SCS7 |
|  | r_0232 | C-22 sterol desaturase (NAD) | CBR1/ERG5/CYB5 or MCR1/ERG5/CYB5 |
|  | r_0441 | **FMN reductase** | **LOT6** |
|  | r_0242 | C-5 sterol desaturase | ERG3 |
|  | r_0259 | ceramide-1 hydroxylase (24C) | SUR2 |
|  | r_0269 | ceramide-4 synthase (24C) | SCS7 |
|  | r_0481 | glutathione oxidoreductase | GRX1/GRX2/ GRX3/GRX4/GLR1 |
|  | r_0806 | nucleoside triphosphatase | YND1 |
|  | **and 29 other reactions** |  |  |
| G (0; 19.8) | r_0439 | ferrocytochrome-c: oxygen oxidoreductase | Cob/COR1QCR7/RIP1/CYC7/ QCR6/QCR9/QCR10/QCR8/ CYT1/QCR2/Cob/QCR7/RIP1 |
|  | r_0773 | NADH:ubiquinone oxidoreductase | NDI1 |
|  | r_0097 | acetolactate synthase | ILV6/ILV2 |
|  | r_0352 | dihydroxy-acid dehydratase (2,3-dihydroxy-3-methylbutanoate) | ILV3 |
|  | r_0096 | acetohydroxy acid isomeroreductase | ILV5 |
|  | r_1022 | succinate-CoA ligase (ADP-forming) | LSC1/LSC2 |
|  | r_0831 | oxoglutarate dehydrogenase (dihydrolipoamide S-succinyltransferase) | KGD1/KGD2/LPD1 |
|  | r_0505 | glycine-cleavage complex (lipoamide) | GCV1/GCV2/GCV3/ LPD1/KGD1/KGD2 |
|  | r_0832 | oxoglutarate dehydrogenase (lipoamide) | KGD1/KGD2/LPD1 |
|  | r_0451 | fumarase | FUM1 |
|  | r_0713 | malate dehydrogenase | MDH1 |
|  | r_0300 | citrate synthase | CIT1/CIT3 |
|  | r_0961 | pyruvate dehydrogenase | PDB1/PDA1/LPD1/PDX1/LAT1 |
|  | r_1021 | **succinate dehydrogenase (ubiquinone-6)** | **SDH2/SDH3/SDH4/YJL045W or SDH1/SDH2/SDH3/SDH4** |
|  | r_0175 | aldehyde dehydrogenase (acetylaldehyde, NADP) | ALD4/ALD5 |
|  | **and 300 other reactions** |  |  |

The proposed deletions of genes involved in glycerol metabolism as a means of improve ethanol production is of particular interest, since several studies have demonstrated that indeed modulation of the activity of this pathway can contribute, at different extents, to improve ethanol production in yeast (reviewed in [26–28]). It can be reasoned that the reduced activity of pathways consuming pyruvate and acetaldeyde can contribute to decrease the diversion of pyruvate, creating a surplus of these metabolites that can then be channeled for ethanol synthesis. However, an eventual link between the metabolism of complex lipids or of glutathione with ethanol production cannot be intuitively established. The fact the set of solutions identified by Momo comprises those expected to be obtained from a biological point of view shows that the modelling is functioning in an appropriate manner.

To validate the results obtained by Momo we considered a reduced set of predictions, among the whole set generated by our software. We eliminated many candidate reactions identified at the Pareto frontier points showing a very small amount of biomass (points F and G) to avoid solutions pointing to increased ethanol production merely by compromising cellular growth. This is a relevant aspect since some of the solutions proposed as means to improve ethanol production could actually be non-viable by compromising cellular viability. We also aimed to include, in the set of strains to be tested, those genes that could not be intuitively be linked to ethanol production. We ended up with a set of 19 different strains to be tested on the capacity of yeast cells to produce ethanol under very high gravity conditions (300g/L glucose as starting sugar concentration). Among the 19 mutant strains tested, four were demonstrated to exhibit increased ethanol levels in comparison with those exhibited by the wild-type strain (14.1% (v/v) *±* 0.5 ethanol for the BY4741 strain): ∆idp1, devoid of the mitochondrial isocitrate dehydrogenase (14.6% *±* 0.6); ∆lot6, devoid of a cytosolic FMN-dependent NAD(P):quinone reductase (16.1% *±* 0.9); ∆ipt1, devoid of a inositolphosphotransferase (14.9% *±* 0.7) and the W303_∆sdh2 mutant devoid of the succinate dehydrogenase subunit Sdh2 (14.9% *±* 0.7, comparing with 14.4% *±* 0.3)(Table 1).

## 4 Discussion and conclusions

The need to optimize more than one feature is a recognized challenge in microbial metabolic engineering. A very common example where it is necessary to deal with many features is the production of compounds, that fall outside of the microbe’s metabolic repertoire, that competes for the carbon source for growth. Although solutions can be envisaged to solve this problem for the bioprocess point of view (using, for example, continuous fermentation reactors where biomass is maintained at a given dilution rate), this is problematic from the modelling point of view since in the standard models only one objective function is used. In the specific case of end-point metabolites, maximization of its production often results in no growth and vice-versa. Other examples where more than one objective has to be envisaged is the need to reduce the synthesis of by-products that could be coupled to the production of the compound of interest but whose total elimination could be difficult or, in some cases, even impossible. This is the case of glycerol, which is a known product of ethanol production undertaken by *S. cerevisiae* required for maintenaince of redox potential and osmobalancing of yeasts cells, thereby rendering its elimination (to prevent carbon diversion) very difficult (as reviewed in [26]).

In this work we present Momo, a framework that is able to consider metabolic networks in multi-objective settings and that also pinpoints for a selected set of reactions that could be deleted in order to reach different combinations of the various objectives selected by the user. Several other tools capable of performing identification of interesting knock-outs have also been developed along time (as reviewed in [6]), out of which OptKnock or OptGene are particularly known [2, 29]. The main difference of Momo in relation with other methods is that it is based on multi-objective otpimization.

One interesting aspect of the Momo approach, in contrast with the bi-level model used by OptKnock, is that in this tool we do not need to impose constraints on the biomass values since all the optimal solutions for different biomass values can be obtained. This is advantageous since it does not require to specify an *a priori* biomass growth rate, which could impact/bias the obtained solution and allows an *a posteriori* analysis of the optimal frontier and choice of the points to enumerate deletions.

In Momo all models are solved using exact methods. This have both the positive aspect that guarantees global optimality and the negative aspect that in practice, depending on the number of objectives to optimize and the choice of *K*, the problem may be too hard to be solved in a reasonable time. Our experiences show that, using a modern laptop, solving instances with *K >* 3 and/or more than 3 objective functions may be prohibitive in practice.

Other advantageous aspects of Momo compared with other approaches is the integration of an enumeration framework that instead of giving one single deletion is able to enumerate candidate(s) that may provide an improved production of the target(s) compound(s). As described above, this approach is based on having a reference flux so we can identify the active reactions to be candidates for the knock-out strategies. This reference flux could come from empirical data or FBA simulations for example. In our case we considered the flux obtained from the FBA model optimizing for biomass. A potential problem may be that this model have many optimal solutions that are different enough to have an impact on the enumeration procedure. An alternative to reduce this impact is to replace the Moma approach by the more computationally costly one known as Room [30] that considers only the presence/absence of flux in a reaction instead of its actual flux value. In our test case discussed in Section 2.4, due to the flux constraints used, the impact of different optimal solutions was not too important in the enumeration (*i.e.* the majority of the deletions enumerated were the same regardless of the FBA solution considered as reference). There are some methods like SimpleOptKnock that also have enumeration features. The drawback with SimpleOptKnock is the fact that it enumerates all possible deletions ranking them trough a score

(wild-type prediction of target production). This results in an unpractical number of deletions to try empirically and often the score is unrelated with the real impact of the deletion. On the other hand, our approach enumerates a reduced number of deletions, instead of all possible ones, thus enabling a more suitable identification of candidates for subsequent experimental approaches.

We have tested the effect of the reaction deletions predicted by Momo for ethanol production, with particular emphasis on reactions whose link with ethanol synthesis was not intuitive and could not be easily anticipated.

Out of the 19 deletions tested, four (deletion of IDP1, IPT1, LOT6 and SDH2) led to improvement in the production titers, comparing with the titers obtained by wild-type cells; these improvements ranging from 2% obtained for the ∆idp1 strain to 13% obtained for the ∆lot6 mutant. Although these improvements might seem small, it is well known that even small increases in the final tier of ethanol represent highly significant add-values to the economy of the bioprocess, highly restrained by the high costs of the distillation step required to obtain ethanol from the fermentation broth [10]. IDP1 and SDH2 encode enzymes that participate in the Krebs cycle and therefore their inactivation may result in a lower flux of the cycle thereby increasing the amount of pyruvate that can then be channeled for increased production of ethanol. In the case of IPT1 and LOT6, which encode respectively a inositolphosphotransferase involved in synthesis of complex sphingolipids and a cytosolic FMN-dependent NAD(P) H:quinone reductase, the improvements that appear to result from their deletions in ethanol production are far less easy to understand. Under the conditions that we have used we could not detect a significant difference between growth of the ∆ipt1 and ∆lot6 strains and of the wild-type strain BY4741 thereby indicating that the improved production exhibited by the mutants does not appear to result from their reduced fitness. Despite the molecular basis underlying the improved production phenotype may not be easily understood, the fact that Momo has identified non-obvious target reactions is of particular high interest.

## 5 Conclusion

In this paper, a new method, called Momo based on a multi-objective mixed integer optimization, was developed and explored with the aim of identifying deletion strategies when several objectives should be taken into account and the variables are both continuous (fluxes) and discrete (deletions). We tested the proposed approach for ethanol and biomass maximization in yeast, identifying all the possible single deletions belonging to the Pareto front. Momo is implemented and freely available at http://momo-sysbio.gforge.inria.fr. It can be easily applied to other cases where several metabolic functions can be optimized.

Ongoing work includes the testing of other objectives to be minimized, such as toxicity, and of other specific linear combinations of reactions to be included in the model. Momo significantly expands the metabolic engineering tools available for identifying the best mutant strains and can therefore provide a wider view of the solution space for improving the production of valuable compounds.

## Supporting information

## Supporting information

Appendix S1 Additional theoretical material on multi-objective optimization.

Table S1 Flux distribution at each Pareto point.

Table S2 All deletion candidates enumerated by Momo.

## Acknowledgements

This work was partially supported by FCT and EraNet in Industrial Biotechnology (6th edition, contract ERA-IB-2-6/0003/2014) and by the European Union, FP 7 BacHBerry (www.bachberry.eu), Project No. FP7-613793. SV acknowledges support by Program Investigador FCT (IF/00653/2012) from FCT, co-funded by the European Social Fund (ESF) through the Operational Program Human Potential (POPH). Funding received by iBB-Institute for Bioengineering and Biosciences (UID/BIO/04565/2013), IDMEC, under LAETA (project UID/EMS/50022/2013), and INESC-ID (UID/CEC/50021/2013), through Fundação para a Ciência e a Tecnologia (FCT) - Portuguese Foundation for Science and Technology, and from Programa Operacional Regional de Lisboa 2020 (Project N. 007317) is also acknowledged. This work was also supported by the São Paulo Research Foundation (FAPESP) grant #2017/05986-2.

## Footnotes

‡ Available at http://polyscip.zib.de/

