## Supplementary material for "Momo – Multi-Objective Metabolic mixed integer Optimization: application to yeast strain engineering"

### Appendix S1

In this section, we present the theoretical background on multi-objective programming and we also discuss a number of efficient methods for solving multi-objective mixed integer (linear) problems.

#### Basic concepts

An optimization problem aims at finding a best solution (optimal solution), among all feasible solutions that satisfy all the constraints of the problem according to the performance of the objective function. When more than one objective or criterion is considered inside an optimization problem, then we are dealing with the concept of multi-objective optimization. In this case, the optimal solution of the optimization problem is not unique.

A general Multi-Objective Mixed Integer Program (MOMIP), with $k$ objective functions and $m$ constraints, is a mathematical optimization problem of the form:

$$\begin{aligned} \max \quad & f(x,y) = (f_1(x,y), \ldots, f_k(x,y)) \\ \text{s.t.} \quad & Ax + Gy \leq b, \\ & x \in \mathbb{Z}_+^n, y \in \mathbb{R}_+^p, \end{aligned} \tag{1}$$

where $f : \mathbb{Z}_+^n \times \mathbb{R}_+^p \longrightarrow \mathbb{R}^k$ is a (multi-criterion) objective function, $A$ is an $m \times n$ matrix, $G$ is an $m \times p$ matrix, and $b \in \mathbb{R}^m$.

We define the set of *feasible solutions* to Problem (1) denoted by $X$ as:

$$X = \big\{(x,y) \in \mathbb{Z}_+^n \times \mathbb{R}_+^p \mid Ax + Gy \leq b\big\}.$$

Observe that $X$ is also called the feasible set in *decision space*. Moreover,

$$Z = \{z \in \mathbb{R}^k \mid z_i = f_i(x,y), (x,y) \in X, i = 1,\dots,k\},$$

denotes the feasible set in *criterion space*.

Let $z^1, z^2 \in \mathbb{R}^k$. We then define the relation $\succ$ as follows:

$$z^1 \succ z^2 \iff z_i^1 \geq z_i^2, i = 1,\dots,k \text{ and } z^1 \neq z^2.$$

Considering the above relation, point $z^2 \in \mathbb{R}^k$ is *dominated* by $z^1 \in \mathbb{R}^k$ if $z^1 \succ z^2$.

The set of *efficient solutions* $X_E$ in decision space is defined as:

$$X_E = \{(x,y) \in X \mid \nexists(\bar{x},\bar{y}) \in X : f(\bar{x},\bar{y}) \succ f(x,y)\}.$$

Set $X_E$ is sometimes called the set of *Pareto-optimal solutions*. All solutions in this set are optimal in the sense that one cannot improve one of the objective functions without worsening one or more of the other objective functions. The set of *non-dominated points* $Z_N$ in the criterion space is given by:

$$Z_N = \{z \in \mathbb{R}^k \mid z = f(x,y), (x,y) \in X_E\}, \tag{2}$$

which is sometimes called the *Pareto-optimal front*.

A non-dominated point $z \in Z_N$ is called *unsupported* if it is dominated by any infeasible convex combination (*i.e.* not belonging to $Z_N$) of other non-dominated points (belonging to $Z_N$); otherwise, it is a *supported* point. A supported non-dominated point $z \in Z_N$ is *extreme* if it is an extreme point of $conv(Z)$; otherwise it is a *non-extreme* point.

These concepts are shown in Figure 1 in the bi-objective case ($k = 2$). Points $A$, $B$,

$C$, $D$, $E$, $F$, and $G$ are non-dominated and point $H$ is dominated. Among non-dominated points, $A$, $B$, $C$, $D$, and $E$ are supported and points $F$ and $G$ are unsupported. Moreover, points $A, B, C$, and $E$ are extreme and point $D$ is non-extreme.

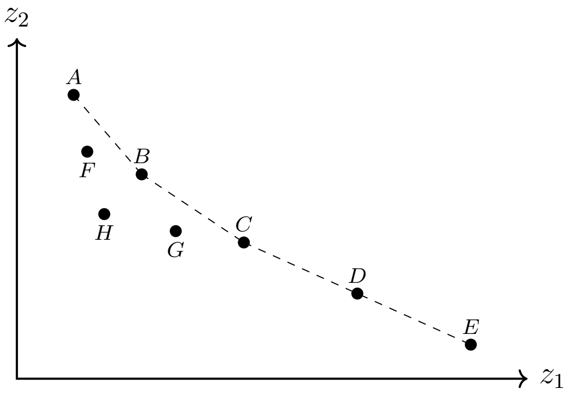


Figure 1: Dominated and non-dominated points in the criterion space.

#### Methods for Multi-Objective Mixed Integer Optimization

A wide range of approaches have been introduced in the literature in order to deal with Multi-Objective Problems (MOPs). However, most of the proposed techniques are developed for continuous multi-objective problems and therefore do not consider the presence of integer decision variables inside the model. More particularly, it is well-known that for multi-objective linear programming problems

$$\max \{Cx : Ax \leq B, x \geq 0\},$$

with $n$ objective functions, the set of efficient solutions is exactly the set of solutions that can be obtained by solving the following single-objective linear problem:

$$\max \Big\{ \sum_{j=1,\ldots,Q} \lambda_j c^j x : Ax \leq B, x \geq 0 \Big\},$$

with $0 < \lambda_j < 1, j = 1, \ldots, n$ and $\sum_{j=1,\ldots,Q} \lambda_j = 1$ [5]. However, this fundamental result is no longer valid in the presence of discrete decision variables in MOMIPs due to the discrete structure of the feasible set. Therefore, there usually exist efficient solutions which are not optimal for any weighted sum of the objectives. Hence, new approaches need to be considered in order to solve MOMIPs efficiently.

We next review the efficient techniques which are developed for solving MOMIPs.

**Weighted Sum Method**

The most traditional and most widely used approach to deal with MOPs/MOMIPs is to scalarize the set of objectives into a single objective by multiplying each objective with a user-supplied weight and then considering the weighted sum of the objectives. Observe that these weights represent the relative importance of each objective and they need to be changed in order to generate different solutions (see [3, 8], for more details).

Although this method is simple and easy to use, it has some disadvantages. As stated earlier, this method may not provide the set of all efficient solutions. Furthermore, two different sets of weight vectors not necessarily lead to two different Pareto-optimal solutions, and it is not clear how to change the weights in order to get an acceptable solution.

**Two-Phase Method**

This method has been the most successful one to solve Bi-Objective Mixed Binary (linear) Programs that was proposed by [12]. It divides the search for non-dominated points into two phases. In Phase 1, the set of supported extreme non-dominated points are usually found by solving a parametric optimization problem. Since supported non-extreme non-dominated points or unsupported non-dominated points may exist, in general it is not possible to find the set of all non-dominated points during Phase 1 of the algorithm. Hence, these points (supported non-extreme and unsupported non-dominated) are found in Phase 2, and finally the set of all Pareto-optimal fronts is identified.

##### Branch-and-Bound Method

The branch-and-bound method has been recognized as one of the most promising and successful methods for solving MOMIPs where some or all of the decision variables are binary. This method aims at finding the set of all non-dominated points by partitioning the problem into disjoint and jointly exhaustive subproblems where a number of branching strategies have been introduced in the literature. Unlike the classical branch-and-bound method that keeps a single incumbent, this method keeps a set of upper bounds of the best solutions found so far. Moreover, the branch-and-bound procedure also manages a lower bound of the subproblems in order to guarantee efficiency of the algorithm.

There exists a number of papers in the literature proposing a branch-and-bound method for solving MOMIPs. For instance [7] developed a sequential generation method for identifying some or all efficient solutions of multiple objective mixed integer linear programming problems, while [6, 9, 10, 11, 13] proposed several branch-and-bound algorithms in order to deal with MOMBPs efficiently.

##### Lifted Weight Space Method

An early version of the lifted weight space algorithm was mentioned in [4]. In order to describe this method, we simplify the presentation of a MOMIP. Given a cost matrix $C \in Q^{k\times(n+m)}$, a constraint matrix $A \in Q^{\ell\times(n+m)}$ and a right-hand side vector $b \in Q^{\ell}$, a MOMIP with $k$ objective functions can be formulated as:

$$\begin{aligned} \min \quad & Cx \\ \text{s.t.} \quad & Ax \leq b \\ & x \in \mathbb{Z}^n \times \mathbb{R}^m. \end{aligned}$$

The algorithm implements the lifted weight space approach in order to compute all feasible solutions which minimize $w^T Cx$ for some $w \geq 0$. More particularly, in each iteration, the algorithm chooses a weight $w = (w_1, \ldots, w_k) \in W = \{w \geq 0 \mid \sum_{i=1}^k w_i = 1\}$ and solves the weighted single-objective problem with cost vector $c = w^T C$. In order to calculate the weights, the algorithm keeps track of the lifted weight space polyhedron $P = \{(a, w) \in \mathbb{R} \times W \mid a \leq w^T Cx, x \in X\}$ where $X$ is the set of solutions found so far. Observe that $P$ is a convex polyhedron and in each iteration, the algorithm chooses a vertex $(a, w) \in P$ such that $w$ has not yet been used in the weighted optimization problem. Every time a new solution is found, $P$ is updated accordingly.

PolySCIP [2] is a solver for multi-objective mixed integer linear programs with an arbitrary number of objectives that utilizes the lifted weight space approach along with the mixed integer linear/non-linear programming solver SCIP [1] to solve the above-mentioned weighted single-objective problems. This solver takes as input a file (mathematical model) which is generated by ZIMPL. ZIMPL is a freely available modelling language to translate the mathematical model of a problem into a mathematical program in the *.mps* file format.
